# Unique properties of Zika NS2B-NS3pro complexes as decoded by experiments and MD simulations

**DOI:** 10.1101/078113

**Authors:** Amrita Roy, Liangzhong Lim, Shagun Srivastava, Jianxing Song

## Abstract

Zika virus can be passed from a pregnant woman to her fetus, thus leading to birth defects including more than microcephaly. It has been recently estimated that one-third of the world population will be infected by Zika, but unfortunately no vaccine or medicine is available so far. Zika NS2B-NS3pro is essential for its replication and thus represents an attractive target for drug discovery/design. Here we characterized conformation, catalysis, inhibition and dynamics of linked and unlinked Zika NS2B-NS3pro complexes by both experiments and MD simulations. The results unveil the unique properties of Zika NS2B-NS3pro which are very different from Dengue one. Particularly, CD and NMR studies indicate that unlike Dengue, the C-terminal region of Zika NS2B with a significant sequence variation is highly disordered in the open conformation. Indeed, MD simulations reveal that up to 100 ns, the Dengue NS2B C-terminus constantly has close contacts with its NS3pro domain. By a sharp contrast, the Zika NS2B C-terminus loses the contacts with its NS3pro domain after 10 ns, further forming a short β-sheet characteristic of the closed conformation at 30 ns. Furthermore, we found that a small molecule, previously identified as an active site inhibitor for other flaviviral NS2B-NS3pro, inhibited Zika NS2B-NS3pro potently in an allosteric manner. Our study provides the first insight into the dynamics of Zika NS2B-NS3pro and further deciphers that it is susceptible to allosteric inhibition, which thus bears critical implications for the future development of therapeutic allosteric inhibitors.

## INTRODUCTION

Zika virus was originally isolated from a sentinel rhesus monkey in the Zika Forest of Uganda in 1947 (1), which is transmitted to humans by Aedes species mosquitoes. Since 2007, large epidemics of Asian genotype Zika virus have been reported around the world (2,3). Recently it has been estimated that one-third of the world population will be infected (4). Most seriously, Zika infection has been found to be associated with serious sequelae such as Guillain-Barré syndrome, and microcephaly in newborn infants of mothers infected with Zika virus during pregnancy (5-8), and consequently WHO has declared a public health emergency for Zika virus (9). Zika virus represents a significant challenge to the public health of the whole world but unfortunately there is no effective vaccine or other therapy available so far.

Zika virus with a single stranded, positive sense RNA genome of 10.7 kb belongs to the flavivirus genus, which also contains Dengue, yellow fever, West Nile, Japanese encephalitis, and tick-borne encephalitis viruses (4,10). Zika virus shares a high degree of sequence and structural homology with other flaviviruses particularly Dengue virus, thus resulting in immunological cross-reactivity (11). Seriously, the current Zika outbreaks are largely localized within dengue-endemic areas, it is thus proposed that preexisting dengue-induced antibodies may enhance Zika infection by antibody-dependent enhancement (ADE), a factor that makes the vaccine approaches extremely challenging (11).

Zika genome is translated into a single ~3,500-residue polyprotein, which is further cleaved into 3 structural proteins and 7 non-structural proteins (10). The correct processing of the polyprotein is essential for replication of all flaviviruses, which requires both host proteases and a viral NS2B-NS3 protease (NS2B-NS3pro) (10,12-18). As a consequence, the flaviviral NS2B-NS3pro has been well-established to be a key target for developing antiviral drugs (10,12-22). In the present study, we successfully obtained two enzymatically active Zika NS2B-NS3pro: one with NS2B and NS3pro linked by extensively used (Gly)_4_-Ser-(Gly)_4_ sequence and another with NS2B and NS3pro unlinked. Subsequently we characterized their conformations, catalysis, inhibition and dynamics by both experiments and molecular dynamics (MD) simulations. The results reveal that despite sharing a similarity to the Dengue one, Zika NS2B-NS3pro does have unique catalytic and inhibitory properties, which appear to result from its high dynamics. Furthermore, we identified the first allosteric inhibitor for Zika NS2B-NS3pro. Taken together, our study provides the first insight into the unique dynamics of the Zika NS2B-NS3pro, which may have key implications in further design of better protease inhibitors for fighting Zika.

## RESULTS

### Cloning, expression and purification of linked and unlinked Zika NS2B-NS3pro

Based on the sequence alignment with NS2B and NS3pro of the Dengue serotype 2 we previously characterized (12), the corresponding Zika sequences were identified for the NS2B (Fig. 1A) and NS3pro (Fig. 1B) of the Asian Zika virus. From synthetic genes with *E. coli* preferred codons, we amplified and subsequently cloned the DNA fragments into His-tagged expression vectors, which encode the isolated NS2B (48-100) with the transmembrane regions deleted (Fig. 1A); as well as isolated NS3 (14–185) (Fig. 1B). The expression level of the recombinant NS2B was extremely low, and consequently we needed to grow many liters of *E. coli* cells to obtain sufficient NS2B for refolding with NS3pro by the same protocol we previously utilized for the Dengue one (12). The refolded NS2B-NS3pro with His-tag (column 2 of Fig. 1C) was cleaved by thrombin covalently linked to beads to remove the His-tag, followed by binding to an excess amount of the Ni^2+^-beads to remove His-tag and uncleaved fusion protein, as well as final purification with FPLC gel-filtration chromatography to obtain unlinked NS2B-NS3pro without His-tag (column 3 of Fig. 1C). We attempted to refold Zika NS3pro without NS2B and found that NS3pro completely precipitated during refolding, thus indicating that Zika NS3pro also absolutely requests its NS2B cofactor for the correct folding, as previously observed on other flavivirus NS3 proteases (12-18).

**FIGURE 1.**
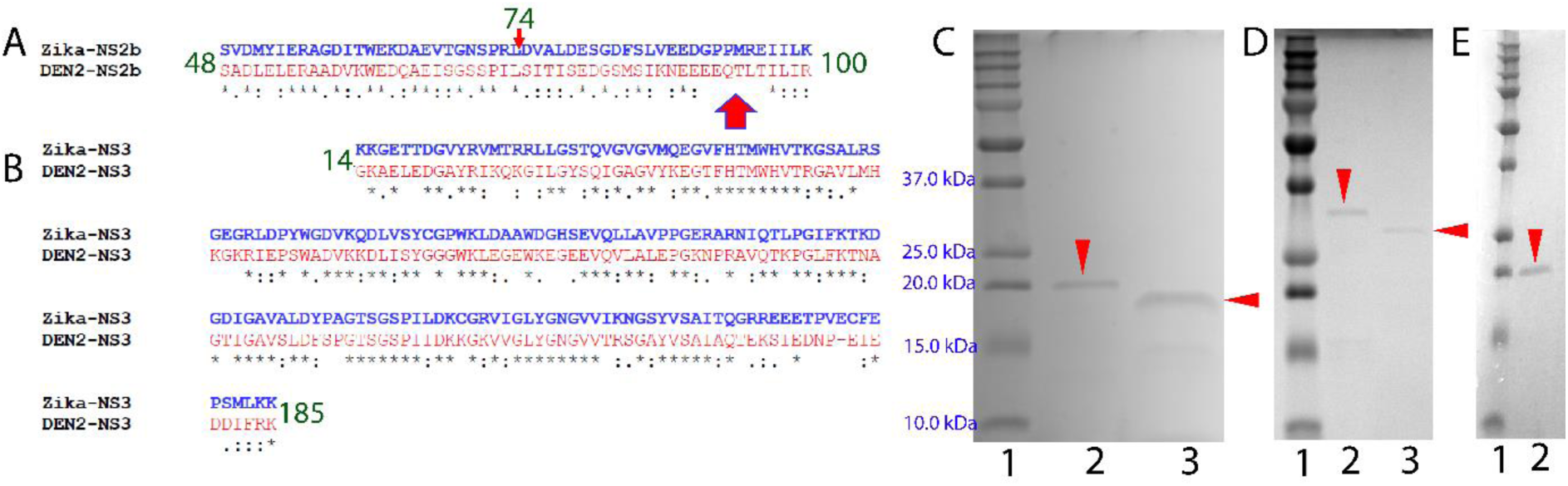
Construction, expression and purification of Zika NS2B-NS3pro complexes. (A) Sequence alignment between NS2B (48-100) of the Dengue and Zika viruses with the transmembrane region removed. The red arrow is used to indicate the region with significant sequence variations. (B) Sequence alignment between NS3pro (14-185) of Dengue and Zika viruses. (C) SDS PAGE of the purification of unlinked Zika NS2B-NS3pro: column 1: molecular weight makers; column 2: unlinked Zika NS2B-NS3pro; column 3: unlinked Zika NS2B-NS3pro with the His-tag removed by the thrombin beads followed by binding to an excess amount of Ni^2+^-beads. (D) SDS PAGE of the purification of linked Zika NS2B-NS3pro: column 1: molecular weight makers; column 2: linked Zika NS2B-NS3pro; column 3: linked Zika NS2B-NS3pro with the His-tag removed by the thrombin beads followed by binding to an excess amount of Ni^2+^-beads. (E) SDS PAGE of the purification of unlinked Zika NS2B(48-74)-NS3pro: column 1: molecular weight makers; column 2: unlinked Zika NS2B(48-74)-NS3pro. Due to the small sizes of NS2B(48-100) and NS2B(48-74), they diffused and thus could not be seen in SDS PAGE.

We also constructed a Zika protease with NS2B and NS3pro linked by a (Gly)4-Ser-(Gly)_4_ sequence extensively used for functional and structural characterization of flavivirus NS2B-NS3pro complexes (13-20). No linked NS2B-NS3pro protein was detected in the supernatant, but we were able to purify a small amount from the inclusion body by the Ni^2+^-affinity chromatography under the denaturing condition, which was confirmed to be the linked NS2B-NS3pro by sequencing with mass spectrometry and western plot with the His-tag antibody. Linked NS2B-NS3pro was successfully refolded by dialysis without any precipitation (column 2 of Fig. 1D). Subsequently, linked NS2B-NS3pro without His-tag (column 3 of Fig. 1D) was obtained by the cleavage with thrombin covalently linked to beads, followed by binding to Ni^2+^-beads and final purification with FPLC gel-filtration chromatography as described above.

### Biophysical characterization

We acquired ^1^H NMR one-dimensional spectra for both linked and unlinked NS2B-NS3pro (Fig. 2A). Both spectra have similar up-field peaks manifested, suggesting that both of them are folded (12,20,23,24). However, it is interesting to note that the peaks of the linked complex are much more broad than those of the unlinked, implying that the linkage may introduce µs-ms conformational dynamics, as previously observed on the linked Dengue NS2B-NS3pro (12,20,23). Indeed, we attempted to acquire ^1^H-^15^N NMR HSQC spectrum of the linked protease, but failed to detect any peaks due to significant peak broadening. By contrast, despite having almost the same sequences, ^15^N-labeled Zika NS3pro in the unlinked complex has a well-dispersed HSQC spectrum, indicating that the NS3pro is well-folded, although many HSQC peaks were still too broad to be detectable (Fig. 2B). Amazingly, ^15^N-labeled Zika NS2B in the unlinked complex has a very narrowly-dispersed HSQC spectrum, in which HSQC peaks of 22 residues are too broad to be detected, and in particular only two out of four Gly residues have HSQC peaks detectable (Fig. 2C). This is very different from what was previously observed on Dengue NS2B in the complex, which showed a well-dispersed HSQC spectrum with peaks of most residues detectable (12,20,23). This implies that Zika NS2B is much more dynamic than Dengue one, in which a portion of residues undergoes µs-ms dynamics, thus becoming too broad to be detectable; while the rest is disordered and absent of tight packing, thus resulting in narrowly-dispersed HSQC peaks (12,23,24).

**FIGURE 2.**
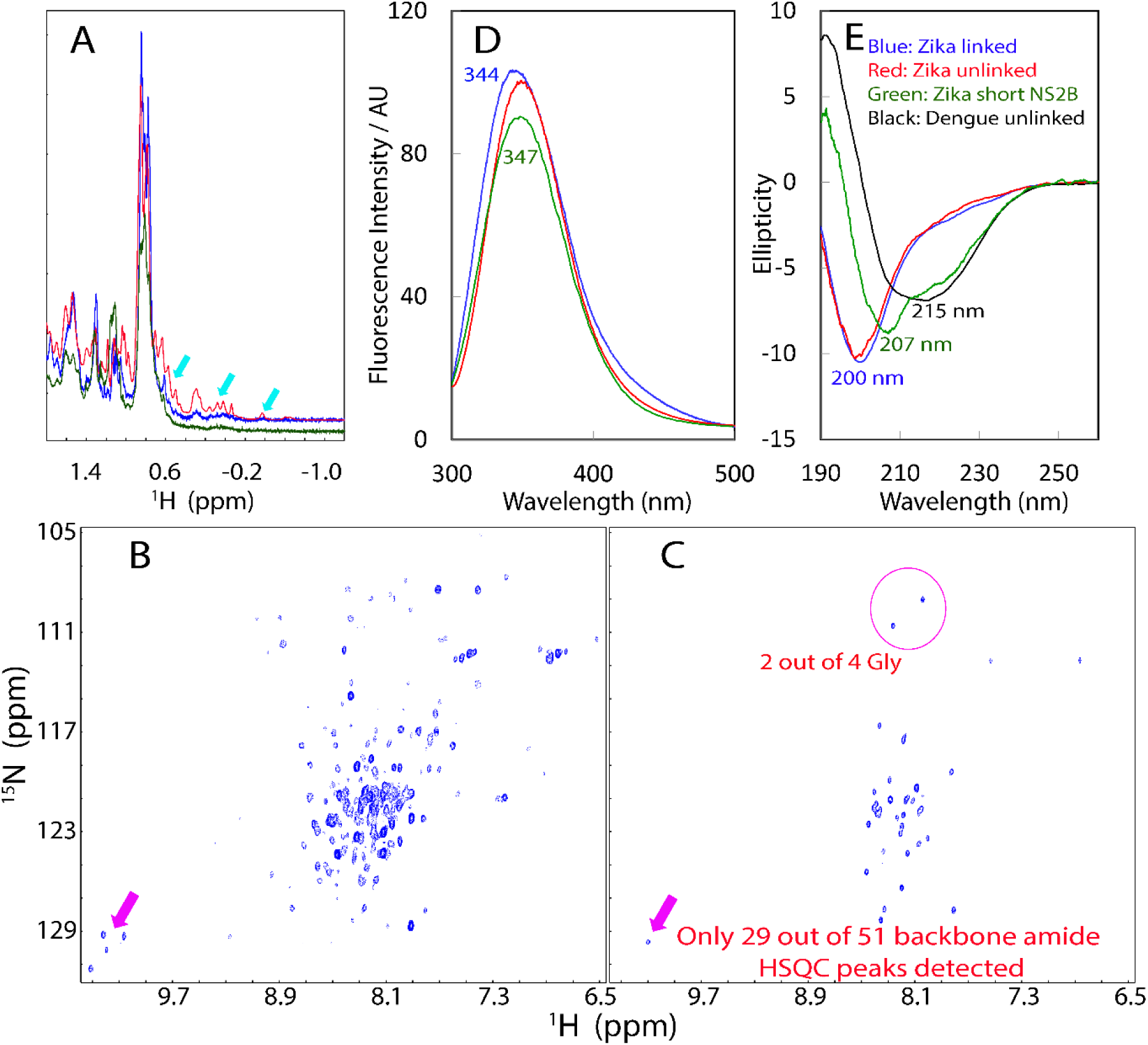
Biophysical characterization of Zika NS2B-NS3pro complexes. (A) One-dimensional ^1^H NMR spectra over -1.2-1.8 ppm of linked (blue) and unlinked (red) Zika NS2B-NS3pro, as well as NS2B(48-74)-NS3pro (green) complexes at a protein concentration of 30 µM. Cyan arrows are used to indicate the very up-field peaks. (B) ^1^H-^15^N HSQC spectrum of ^15^N-labeled NS3pro in complex with unlabeled NS2B at a protein concentration of 30 µM. (C) ^1^H-^15^N HSQC spectrum of ^15^N-labeled NS2B in complex with unlabeled NS3pro at a protein concentration of 30 µM. Pink arrows are used to indicate the HSQC peaks of Trp side chains in NS3pro and one in NS2B. (D) Emission spectra of the intrinsic UV fluorescence of linked (blue) and unlinked (red) Zika NS2B-NS3pro, as well as NS2B(48-74)-NS3pro (green) complexes at a protein concentration of 10 µM. (E) Far-UV CD spectra of linked (blue) and unlinked (red) Zika NS2B-NS3pro, as well as NS2B(48-74)-NS3pro (green) complexes at a protein concentration of 10 µM, together with that of Dengue NS2B-NS3pro complex previously obtained (black) (12).

We also collected spectra of the intrinsic UV fluorescence from Trp residues for both linked and unlinked NS2B-NS3pro complexes (Fig. 2D), both of which have similar spectra with the emission maxima at ~344 nm, very similar to what were observed on the NS2B-NS3pro complexes of all four Dengue serotypes (348 nm) (16). This suggests that in Zika NS2B-NS3pro, all 4 Trp residues are similarly buried as Dengue ones, as Trp residue in the unfolded proteins has an emission maximum wavelength > 352 nm (24). Strikingly, however, both linked and unlinked Zika NS2B-NS3pro complexes have far-UV CD spectra with the maximal negative signal at the wavelength of ~201 nm and is lacking of any positive signal below 200 nm, which is very different from that of unlinked Dengue one of the same length (12), which has the maximal negative signal at 217 nm and large positive signal at 192 nm (Fig. 2E). This strongly suggests that Zika NS2B-NS3pro complexes contain much more disordered regions than the Dengue, completely consistent with the NMR HSQC result of the ^15^N-labeled NS2B (Fig. 2C). Unfortunately, the low solubility and µs-ms conformational dynamics prevented us from conducting NMR assignments of Zika NS2B-NS3pro by triple-resonance NMR experiments.

### Characterization of the enzymatic catalysis

We subsequently characterized the enzymatic properties of both linked and unlinked Zika NS2B-NS3pro complexes. First, we found that the activity of both complexes showed a very similar dependence on the pH values, with the optimal pH at ~9.5 (Fig. 3A), very similar to other flavivirus NS2B-NS3pro linked by (Gly_4_)-Ser-(Gly_4_) (14-18). Amazingly, this is very different from Dengue NS2B-NS3pro, for which the optimal pH significantly switched from basic pH to neutral pH upon separating NS2B from NS3pro (23). Second, the catalytic activity reduced significantly upon increasing the concentrations of NaCl (Fig. 3B). In the presence of 150 mM NaCl, the catalytic activity is only 13.6% and 8.2% respectively of the linked and unlinked enzymes in 50 mM Tris buffer at pH 8.5.

**FIGURE 3.**
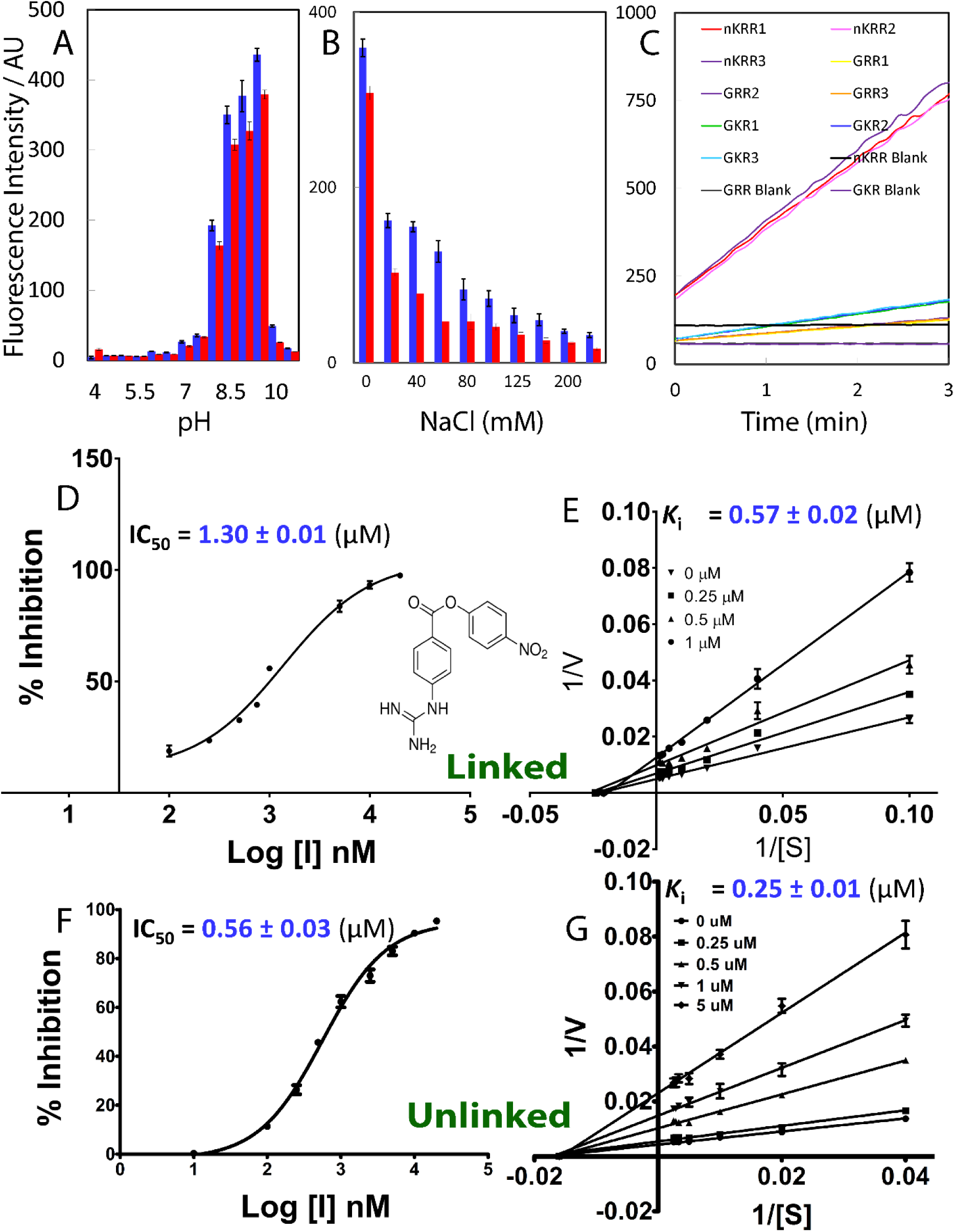
Characterization of Enzymatic catalysis and inhibition. (A) Enzymatic activities of linked (blue) and unlinked Zika NS2B-NS3pro complexes at different pH values. (B) Enzymatic activities of linked (blue) and unlinked (red) Zika NS2B-NS3pro complexes in 50 mM Tris buffer at pH 8.5 with additional addition of NaCl at 0, 20, 40, 60, 80, 100,125,150, 200, 250 mM. (C) The tracings of fluorescence intensity within 3 min for three different substrates cleaved by linked Zika NS2B-NS3pro complex: Bz-nKRR-AMC, Boc-GRR-AMC and Boc-GKR-AMC; as well as three assay buffers without the protease. Fluorescence intensity is reported in arbitrary units. (D) Fitting curve of IC_50_ for linked Zika NS2B-NS3pro inhibited by a small molecule inhibitor p-Nitrophenyl-p-guanidino benzoate. (E) The Lineweaver-Burk plot for linked Zika NS2B-NS3pro. (F) Fitting curve of IC_50_ for unlinked Zika NS2B-NS3pro. (E) The Lineweaver-Burk plot for unlinked Zika NS2B-NS3pro. [S] is the substrate concentration; v is the initial reaction rate. Both curves were generated by the program GraphPad Prism 7.0.

Furthermore, to allow comparison with the kinetic results previously published on profiling the substrate specificity for the (Gly)_4_-Ser-(Gly)_4_ linked NS2B-NS3 proteases of all four Dengue serotypes (17), here we selected the same three substrates, which were also extensively used for other flavivirus NS2B-NS3pro, namely Bz-nKRR-AMC; Boc-GRR-AMC and Boc-GKR-AMC. We also measured the proteolytic activities of our linked and unlinked Zika NS2B-NS3pro complexes in the same buffer as previously published (17). Interestingly, as shown in Fig. 3C, the Zika NS2B-NS3pro efficiently cleaved Bz-nKRR-AMC, but unexpectedly showed very weak activity on other two. This implies that in addition to requesting dibasic residues at the P1 and P2 sites characteristic of the flaviviral NS3 proteases, Zika NS2B-NS3pro appears to have the higher request of a basic residue at P3 site than the Dengue complexes (17). However, as our current focus was not on profiling substrate specificity, in the present study, we did not test on more substrates but measured kinetic constants of linked and unlinked Zika complexes on Bz-nKRR-AMC.

As shown in Table 1, linked and unlinked Zika complexes only have slightly different kinetic constants. On the other hand, the linked Zika complex showed much larger differences from those of Dengue complexes (17). For example, linked Zika complex has a Km of 43.38 µM, which is ~7 folds of the Dengue 1 (6.2 µM); ~3.6 folds of the Dengue 2 and Dengue 3 (12 µM); and ~6 folds of the Dengue 4 (7.3 µM) on the same substrate. This implies that the Zika complex has the lower affinity to this substrate than all four Dengue ones. However, the linked Zika NS2B-NS3pro has a kcat (48.89 S^-1^) much larger than those of all four Dengue (0.32, 1.4, 0.61 and 2.8 S^-1^ respectively). Consequently, the catalytic efficiency as reflected by kcat/Km (1,128,681 M^-1^S^-1^) is ~3 folds of that of Dengue 4 (380,000 M^-1^S^-1^), the largest among the four Dengue complexes (17).

**Table 1.** Kinetic parameters of catalysis and inhibition of linked and unlinked Zika NS2B-NS3pro complexes.

|  | **Linked** | **Unlinked** |
| --- | --- | --- |
| **K<sub>m</sub>** (μM) | 43.38 ± 1.99 | 60.22 ± 0.66 |
| **k<sub>cat</sub>** (S<sup>-1</sup>) | 48.89 ± 0.41 | 93.10 ± 1.10 |
| **k<sub>cat</sub>/K<sub>m</sub>** (M<sup>-1</sup>S<sup>-1</sup>) | 1,128,681 ± 56,053 | 1,545,992 ± 20,045 |
| **V<sub>max</sub>** (μM//min) | 205.20 ± 1.60 | 318.37 ± 3.70 |
| **IC<sub>50</sub>** (μM) | 1.30 ± 0.01 | 0.56 ± 0.03 |
| **K<sub>i</sub>** (μM) | 0.57 ± 0.02 | 0.48 ± 0.03 |

Interestingly, the kinetic constants were just published for a smaller Zika NS2B-NS3pro which was also linked by the same (Gly)_4_-Ser-(Gly)_4_ but had almost all disordered regions removed (25). It has a Km of 18.3 µM and kcat of 44.6 S^-1^ respectively. While the kcat values are very similar, the Km value is ~2.4-fold less than our current one. This small difference is likely due to: 1) the difference of substrate as Bz-nKKR-AMC was used in the study (25); or/and 2) the presence of more unstructured regions in our construct, 3) or/and the difference of the buffers: our buffer is 50 mM Tris pH 8.5 while the buffer in the publication (25) is only 10 mM Tris pH 8.5. It is well-established that high salt concentrations reduced the activity of the flaviviral proteases (12-18). Indeed, we measured the activity of our linked Zika protease in 10 mM Tris pH 8.5, and the catalytic activity is 1.36-time higher.

### Characterization of the enzymatic inhibition

Previously, a small molecule p-Nitrophenyl-p-guanidino benzoate (Fig. 3D) has been identified to specifically bind to the active site of West Nile NS2B-NS3pro by molecular dynamics (MD) simulations, followed by NMR binding confirmation (19). Later, this molecule has been extensively demonstrated to bind Dengue NS2B-NS3pro in a similar manner by NMR (12,20,23). Here we have measured its inhibitory effect on both linked and unlinked Zika NS2B-NS3pro complexes. Remarkably, it showed strong inhibitory activities, with IC_50_ values of 1.30 ± 0.01 (Fig. 3D) and 0.56 ± 0.03 (Fig. 3F) µM; and *K*i values of 0.57 ± 0.02 (Fig. 3E) and 0.48 ± 0.03 (Fig. 3G) µM respectively for linked and unlinked Zika complex (Table 1). Most strikingly, as clearly evidenced from Lineweaver-Burk plots for both linked (Fig. 3E) and unlinked (Fig. 3G) Zika complexes, the small molecule acts as a non-competitive inhibitor for both forms. We have searched all literatures citing the original paper (19), but only found IC_50_ value of this inhibitor on West Nile NS2B-NS3pro complex (34.2 µM) in the original paper (19). Therefore, this inhibitor could inhibit the enzymatic activity of Zika NS2B-NS3pro much more potently than that of West Nile complex. While it remains unclear whether this molecule inhibits other flaviviral NS2B-NS3pro complexes in a competitive or non-competitive manner, our present results clearly reveal that it inhibits Zika NS2B-NS3pro complexes in a non-competitive manner, fundamentally different from a just reported peptidomimetic boronic acid inhibitor which competitively inhibited Zika NS2B-NS3pro (25). Our results thus decoded for the first time that Zika NS2B-NS3pro in fact has a pocket, to which small molecules can bind to trigger allosteric inhibition of the enzymatic activity.

### Dynamic behaviors as revealed by MD simulations

Low solubility and high dynamics of our longer Zika NS2B-NS3pro constructs prevented from further determination of its structure by crystallography or NMR. On the other hand, Zika and Dengue viruses have a high sequence identity for the NS2B (35.19%) and NS3pro (56.67%) respectively, which is sufficient to allow to build up a homology model (26,27). Therefore, with MODELLER software suite (9v10) (26), we generated a homology model of the Zika NS2B-NS3pro in the “open” form by use of a Dengue crystal structure (PDB code: 2FOM) as a template. This Dengue crystal structure represents the “open” form, which is adopted by the flaviviral NS2B-NS3pro complexes in the absence of any inhibitors (12,18,21,22). The obtained Zika model only has 0.13 Å of root-mean-square-distance (RMSD) of heavy atoms from the Dengue complex.

Molecular dynamics (MD) simulation represents a powerful tool to gain insights into roles of protein dynamics in the enzymatic catalysis (28), and we have previously utilized it to study the Dengue NS2B-NS3pro in the “open” form (12), as well as the dynamically-driven allosteric mechanisms of the SARS 3C-like protease, which also shares the chymotrypsin fold to host the catalytic machinery (29,30). Here with the exact same protocols we previously used for the Dengue (12), we conducted MD simulations up to 100 ns for Zika and Dengue NS2B-NS3pro complexes in both open and closed forms, with each having four independent simulations.

Fig 4A and 4B show the root-mean-square deviations (RMSD) trajectories averaged over four independent MD simulations respectively for NS3pro and NS2B of both Zika and Dengue complexes in the open form. While the Zika NS2B and NS3pro have the averaged RMSD values of 5.33 ± 0.45 and 1.73 ± 0.12 Å respectively, the Dengue NS2B and NS3pro have 4.23 ± 0.46 and 1.61 ± 0.11 Å respectively. As seen in Fig. 4A, Zika NS3pro only has a slightly higher RMSD than Dengue one and interestingly, after 60 ns their average RMSD trajectories become similar. By contrast, Zika NS2B has a constantly larger average RMSD trajectory than Dengue one during the whole 100-ns simulations. This indicates that both Zika NS2B and NS3pro have intrinsic dynamics higher than those of the Dengue, and particularly the Zika NS2B is much more dynamic than the Dengue one.

**FIGURE 4.**
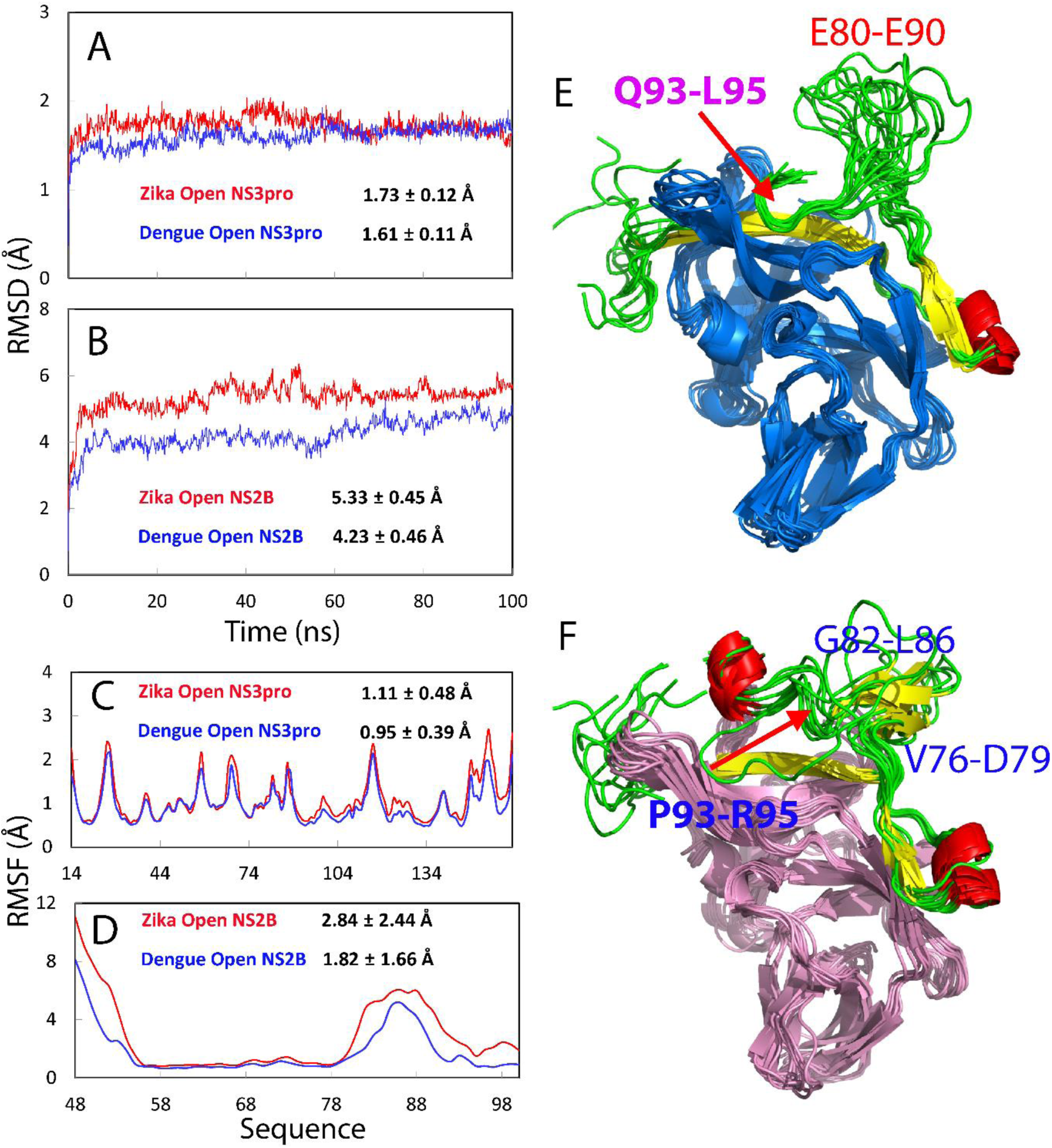
Dynamics of Zika and Dengue NS2B-NS3pro complexes in open forms by MD simulations. RMSD trajectories of the Cα atoms (from their positions in the energy minimized structures) averaged over four independent MD simulations for NS3pro (A) and NS2B (B) in the Zika (red) and Dengue (blue) protease complexes in the open conformations. RMSF of the Cα atoms averaged over four independent MD simulations for NS3pro (C) and NS2B (D) in the Zika (red) and Dengue (blue) protease complexes in the open conformation. Structure snapshots (one structure for 10-ns interval) of MD simulations for Dengue (E) and Zika (F) NS2B-NS3pro complexes in the open conformations. NS2B was specially colored as α-helix (red), β-sheet (yellow) and loop (green).

Furthermore, although the overall patterns of the root-mean-square fluctuations (RMSF) are similar for NS3pro (Fig 4C) and NS2B (Fig 4D) of Zika and Dengue complexes, the RMSF values averaged over four simulations of the Zika ones are all larger than those of the Dengue. In particular, the Zika NS2B has much larger RMSF over the N-terminal 6 residues C-terminal 22 residues (Fig 4D). A close inspection of the structure snapshots reveals that although the Dengue NS2B residues Glu80-Glu90 have large conformational fluctuations, the residues Gln93-Thr94-Leu95 constantly have close contacts with the NS3pro residues Leu31-Gly32-Tyr33-Ser34 in the whole 100-ns simulations (Fig 4E), and consequently have small RMRF over this short fragment (Fig 4D). By a sharp contrast, the Zika residues Pro93-Met94-Arg95 corresponding to Dengue Gln93-Thr94-Leu95 lose the contact with NS3pro after 10 ns and consequently the Zika NS2B C-region becomes highly exposed (Fig. 4F).

Fig. 5A shows the structure snapshots of Dengue NS2B, which highlights the very similar conformations of Gln93-Leu95 at different simulation time points. Most strikingly, the structure snapshots of Zika NS2B (Fig 5B) reveal that only after 30 ns, a short β-sheet is formed with two β-strands over residues Val76-Asp79 and Gly82-L86, which is further illustrated by the structure snapshots of Zika NS2B-NS3pro complex (Fig. 5C). This is a significant finding as in the closed conformation, this β-sheet further get packed to NS3pro, even with residues Ser80-Gly81-Asp82 directly contacting the inhibitor (25). Unfortunately, the formation of such packing is expected to occur at least over µs-ms time scale, which still remains challenging to be simulated by MD.

**FIGURE 5.**
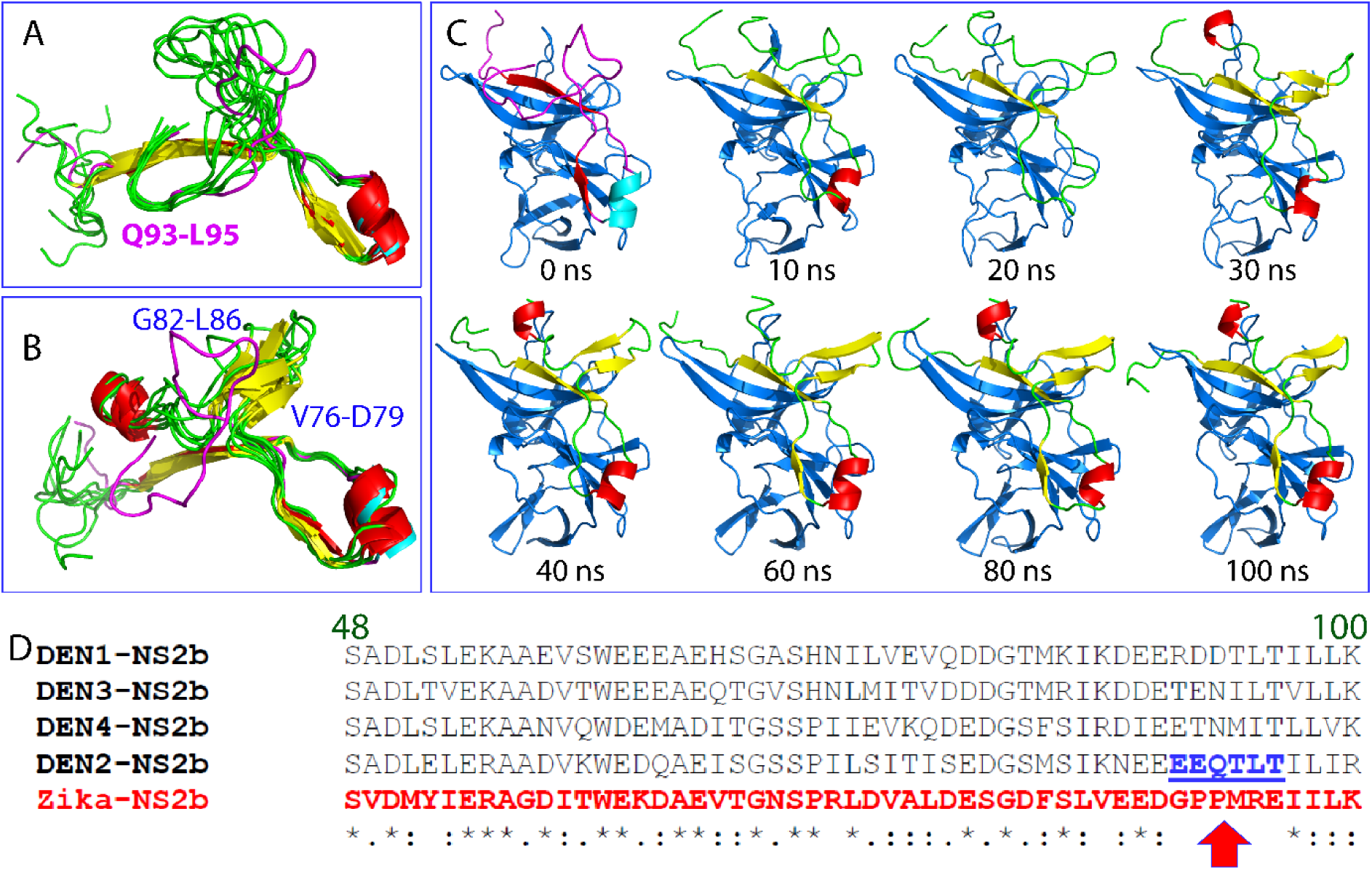
Significant difference of dynamics between Zika and Dengue NS2B in the open conformation by MD simulations. (A) Structure snapshots of NS2B in Dengue NS2B-NS3pro complex in the open conformation in MD simulations at 0, 10, 20, 30, 40, 60, 80, 100 ns. (B) Structure snapshots of NS2B in Zika NS2B-NS3pro complex in the open conformation in MD simulations at 0, 10, 20, 30, 40, 60, 80, 100 ns. (C) Structure snapshots of Zika NS2B-NS3pro complex in the open conformation in MD simulations at 0, 10, 20, 30, 40, 60, 80, 100 ns. NS2B was specially colored as α-helix (cyan), β-sheet (red) and loop (pink) for the one at 0 ns and α-helix (red), β-sheet (yellow) and loop (green) for the rest. (D) Sequence alignment of NS2B (48-100) of Zika and four serotype Dengue viruses. The red arrow is used to indicate the region with significant sequence variations.

So why do Zika and Dengue NS2B have such radical differences in conformations and dynamics? Examination of NS2B sequences reveals that the Zika NS2B region corresponding to the Dengue NS2B Glu91-Thr96 shows no sequence homology at all (Fig. 5D), thus implying that the sequence variations over this region may mainly account for the loss of the contact between the Zika NS2B and NS3pro corresponding to the Dengue NS2B Gln93-Leu95 and NS3pro Leu31-Ser34, thus leading to the high dynamics of Zika NS2B C-region, which is successfully detected by CD and NMR spectroscopy, as well as pinpointed by MD simulations. We also attempted to assess the dynamics of Zika and Dengue NS2B-NS3pro complexes in the closed conformation. To achieve this, we performed MD simulations up to 100 ns on the crystal structures of Zika (5LC0) (25) and Dengue (3U1I) (31) NS2B-NS3pro complexes with the small molecule inhibitors removed. We remove the small molecule inhibitors because: 1) we aimed to find whether the removal of the small molecules would trigger a difference in dynamic instability for Zika and Dengue complexes. 2) Currently the force-field parameters for simulating small molecules are still not universally established.

Fig 6A and 6B show the root-mean-square deviations (RMSD) trajectories averaged over four independent MD simulations respectively for NS3pro and NS2B of both Zika and Dengue complexes in the closed form. Interestingly, for the closed conformation without inhibitors, Dengue complex is more dynamic than the Zika one. While the Dengue NS3pro has the averaged RMSD values of 2.12 ± 0.16 Å, slightly higher than that of Zika (1.96 ± 0.22 Å), the Dengue NS2B has the averaged RMSD values of 8.01 ± 0.82 Å, much higher than that of Zika (6.82 ± 0.78 Å). Furthermore, although the overall patterns of the root-mean-square fluctuations (RMSF) are similar for NS3pro (Fig 6C) and NS2B (Fig 6D) of Zika and Dengue complexes, the RMSF values averaged over four simulations of the Dengue ones are all larger than those of the Zika. In particular, the Dengue NS2B has relatively larger RMSF than Zika one over the C-terminal 7 residues (Fig 6D). It is very interesting to observe here that although the majority of contacts of the small molecule inhibitors are with NS3pro, their removal triggers much more significant changes of dynamics of NS2B than those of NS3pro for both Zika and Dengue complexes. Again, as the current simulations only reached 100 ns, the short β-sheet still persists and remains packed with NS3pro for both Dengue (Fig. 6E) and Zika (Fig. 6F) complexes.

**FIGURE 6.**
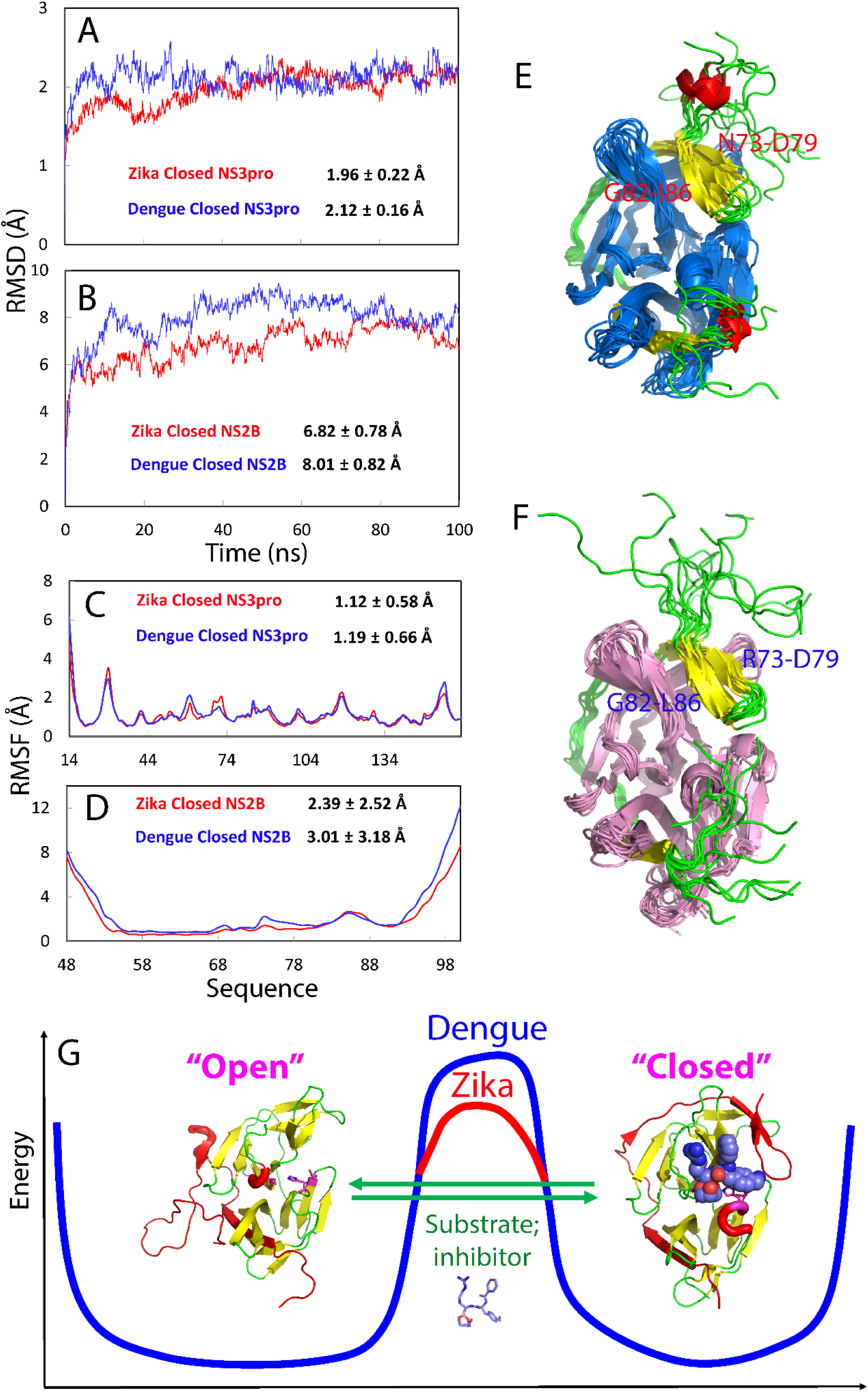
Dynamics of Zika and Dengue NS2B-NS3pro complexes in the closed form by MD simulations. RMSD trajectories of the Cα atoms (from their positions in the energy minimized structures) averaged over four independent MD simulations for NS3pro (A) and NS2B (B) in the Zika (red) and Dengue (blue) protease complexes in the closed conformation. RMSF of the Cα atoms averaged over four independent MD simulations for NS3pro (C) and NS2B (D) in the Zika (red) and Dengue (blue) protease complexes in the closed conformation. Structure snapshots (one structure for 10-ns interval) of MD simulations for Dengue (E) and Zika (F) NS2B-NS3pro complexes in the closed conformation. NS2B was specially colored as α-helix (red), β-sheet (yellow) and loop (green). (G) Proposed diagram showing the conformational exchange between the “open” and “closed” forms of the NS2B-NS3pro complex, which is separated by energy barriers of different heights for Zika and Dengue. The spheres are used to indicate the inhibitor molecule.

### Characterization of Zika NS2B(48-74)-NS3pro complex

To test the simulation results showing that the high dynamics of Zika NS2B-NS3pro complex detected by CD and NMR is mostly resulting from the C-terminal region of Zika NS2B, we cloned and expressed a His-tagged Zika NS2B(48-74) with the C-region deleted. Despite its low expression level and insolubility, by growing many liters of *E. coli* cells, we managed to purify a sufficient amount of NS2B(48-74) for refolding with NS3pro. Interestingly, NS2B(48-74) is capable of forming a soluble complex with NS3pro (Fig. 1E). However, its NMR peaks are much more broad even than those of linked NS2B-NS3pro (Fig. 2A), most likely due to µs-ms conformational dynamics or/and dynamic aggregation. On the other hand, its intrinsic UV fluorescence spectrum indicates that its four Trp residues are similarly buried as linked and unlinked Zika complexes with the full-length NS2B (Fig. 2D). Most interestingly, as judged from its CD spectrum which has the maximal negative signal shifted to 207 nm and has positive signal at 191 nm (Fig. 2E), this complex with the C-region of NS2B deleted indeed contains the less amount of disordered region than both linked and unlinked Zika complexes with the full-length NS2B (Fig. 2E). However, despite being less disordered, this complex showed no detectable enzymatic activity even with the protease concentration up to 20 µM, suggesting that this complex is completely inactive. Previously, we have generated a truncated Dengue NS2B with the C-terminal residues deleted but the short NS2B lost the capacity in forming a soluble complex with its NS3pro. However, we generated another truncated NS2B with only residues 77-84 deleted, which was designated as NS2B(48-100; Δ77–84). Interestingly, NS2B(48-100; Δ77–84) was able to form a soluble complex with its NS3pro, which appeared to be highly disordered as reflected by its CD spectrum, as well as highly dynamic as judged by its NMR spectrum (12).

Together, these results clearly indicate that unlike Dengue one, Zika NS2B indeed has a dynamic and disordered C-terminal region even in the enzymatically active complexes with its NS3pro domain. Nevertheless, our results suggest that this C-terminal region is absolutely required for implementing the catalytic actions, implying that the closed conformation might be indeed the enzymatically-active conformation, as previously speculated (20-23,31), because the deletion of the C-terminal region completely abolishes the potential of Zika NS2B-NS3pro complex to form the closed conformation. Additionally, despite being highly disordered, the C-region might have other functional roles such as to maintain functional dynamics to facilitate HCV replication (32,33).

## DISCUSSION

Knowledge of catalysis, structures and dynamics of all conformational states is beneficial to design of inhibitors of high affinity and specificity for enzymes including viral proteases (28). This is particular relevant to flaviviral NS2B-NS3 proteases which appear to need a transition from the open (inactive) to closed (active) conformations to achieve their catalytic actions (20-23,31). As well demonstrated (18,21-23,25,31), the “open” (inactive) and “closed” (active) forms of the flaviviral NS2B-NS3 proteases have very similar structures of the catalytic domains (NS3pro) but differ mostly in the structures and dynamics of NS2B: the “open” is characteristic of the highly unstructured and dynamic C-half of NS2B, while the “closed” has the C-half adopting a well-folded β-hairpin which become tightly contacted with the NS3pro chymotrypsin fold (Fig. 6G). As a consequence, better understanding of their conformational dynamics may provide key clues to design inhibitors by blocking the conversion from the open to closed conformations.

Here, we have successfully obtained both linked and unlinked Zika NS2B-NS3pro complexes, which, despite sharing similarity, do have some unique catalytic features including the lower substrate affinity but higher efficiency than those of the complexes of all four Dengue serotypes. Most strikingly, as reported by CD and NMR HSQC spectroscopy, Zika NS2B appears to be much more dynamic and disordered than Dengue NS2B of the same length as we previously studied (12). Indeed, in preparation of our current manuscript, a crystal structure was published on Zika NS2B-NS3pro which was linked with (Gly)_4_-Ser-(Gly)_4_ but had almost all flexible regions of both NS2B and NS3pro deleted (25). Interestingly, it was reported that unlike Dengue complexes whose high-quality crystals could be obtained for the “open” form, this small Zika complex could only be crystallized in complex with a tight competitive inhibitor, and consequently the structure represents a “closed” form (Fig. 6), in which the characteristic β-sheet is formed over Arg73-Asp79 and Gly82-Leu86 of the NS2B C-half and further becomes tightly packed against NS3pro; and even has residues Ser80-Gly81-Asp82 directly contacting the inhibitor (Fig. 6G).

To obtain high resolution insights into the dynamic behaviors, we have performed 100-ns MD simulations for Zika and Dengue NS2B-NS3pro complexes in both open and closed conformations. The results reveal that while the Dengue NS2B C-terminal residues Gln93-Leu95 constantly have close contacts with the NS3pro domain in all four independent trajectories of 100-ns simulations of the open conformation (Fig. 4E), the Zika NS2B C-terminal residues lose the close contacts with its NS3pro domain and become highly exposed to bulk solvent only after 10 ns. This is completely consistent with NMR HSQC spectrum of ^15^N-labeled Zika NS2B in complex with unlabeled NS3pro (Fig. 2C), indicating that at least half of the Zika NS2B residues are highly dynamic. This unusually high dynamics appear to result mainly from the large sequence difference over the NS2B C-terminal residues 91-96 between Zika and Dengue (Fig. 5D). Indeed, a short Zika NS2B with residues 75-100 deleted is sufficient to form a complex with its NS3pro, which has the less disordered structure but is enzymatically inactive, thus not only confirming that the Zika NS2B C-terminal residues is highly disordered, but also supporting the previous proposal that the formation of the closed conformation is essential for the enzymatic catalysis.

Amazingly, a short β-sheet characteristic of the closed conformation is formed only at 30 ns of simulations of Zika NS2B-NS3pro in the open conformation (Fig. 5). Furthermore, the contact between Dengue NS2B Gln93-Leu95 and its NS3pro only existing in the “open” form is completely incompatible with the structure of the NS2B C-half in the “closed” conformation (Fig. 6E). As such, the loss of this contact in Zika NS2B-NS3pro in the open conformation implies that the energy barrier separating the “open” and “closed” forms of the Zika complex might be lower than that of the Dengue complex (Fig. 6G). In other words, according to “conformational selection” theory (34), the lower energy barrier leads to the more profound conformational exchanges between the “open” and “closed” forms of the Zika complex even in the free state.

Our study also reveals that a small molecule previously established as an active site inhibitor is able to inhibit both linked and unlinked Zika NS2B-NS3pro complexes at high affinity. Most importantly, this molecule acts as an allosteric inhibitor for both Zika NS2B-NS3pro complexes, fundamentally different from a recently reported peptidomimetic boronic acid inhibitor which is a competitive inhibitor (25). This suggests that Zika NS2B-NS3pro is susceptible to allosteric inhibition. Furthermore, as this allosteric inhibitor also has a structure scaffold very different from the peptidomimetic boronic acid inhibitor, our discovery might open up a new avenue for the future development of allosteric inhibitors, which is highly demanded to achieve therapeutic inhibition of flaviviral NS2B-NS3pro complexes (35). Currently we are focused on optimizing various conditions which would allow atomic resolution studies on the structures, dynamics and inhibitor design for Zika NS2B-NS3pro by NMR spectroscopy (12,19,20,23,36-38).

## EXPERIMENTAL PROCEDURES

### Plasmid construction

The identified genes encoding NS3pro (1-185) and NS2B (1-130) from the Asian Zika strain 8375 (GenBank ID: KU501217.1) were optimized and synthesized by GenScript (Piscataway, NJ). With designed primers, the genes by GenScript were used as templates for amplifying DNA fragments encoding the isolated NS3 (14–185) and NS2B (48–100) with the transmembrane regions removed, as well as NS2B (48–74) with the C-region further deleted. Furthermore, DNA fragments were designed for linking NS2B (48-100) to NS3 (14–185) by (Gly)_4_-Ser-(Gly)_4_ using overlap PCR as previously described (14). Amplified DNA fragments were subsequently cloned into His-tagged pET28a vector (Novagen). DNA sequences of all constructs were verified by automated DNA sequencing.

### Protein expression and purification

All pET28a vectors containing different Zika genes were transformed into *Escherichia coli* BL21 (DE3) Star cell, which was cultured in Luria-Bertani broth containing 25 μg/ml kanamycin at 37 °C until the A600 reached 0.6. Protein expression was induced with 1 mM isopropyl β-D-thiogalactopyranoside (IPTG) for 4 h at 37 °C. The cell pellets were resuspended in cold PBS buffer at pH 7.4, containing 10 mM β-mercaptoethanol and 8M urea and the supernatant containing the recombinant proteins were purified by Ni-NTA affinity column under denaturing condition. Eluted fractions was subjected to dialysis against PBS pH 7.4, 10 mM β-mercaptoethanol buffer at 4 °C overnight to allow the refolding of the protease. The refolded protease was subjected to thrombin cleavage using thrombin-agarose beads from Thrombin CleanCleave^TM^ Kit (Sigma-aldrich, St. Louis, MO), followed by binding to an excess amount of Ni-NTA beads to remove His-tag and uncleaved fusion protein, as well as a FPLC purification on a gel filtration column (HiLoad 16/60 Superdex 200). Recombinant protease samples were checked by SDS-PAGE, molecular weights verification with ESI-MS and protein sequencing with time-of-flight-mass spectrometer (Applied Biosystems). Protein concentration was determined by the UV spectroscopic method with 8 M urea (12).

### Fluorescence, CD and NMR experiments

Intrinsic UV fluorescence spectra were measured with a Cary Eclipse fluorescence spectrophotometer as we previously described (24) with the excitation wavelength at 280 nm. Circular dichroism (CD) experiments were performed on a Jasco J-1500 spectropolarimeter and data from five independent scans were added and averaged (12,24). All NMR experiments were acquired on an 800 MHz Bruker Avance spectrometer equipped with pulse field gradient units as described previously (12,24,36).

### Enzymatic Activity and Kinetics

To allow comparison with the enzymatic kinetic parameters reported on the NS2B-NS3pro complexes of four Dengue serotypes (17), we selected three fluorophore-tagged substrates previously used (17): namely Bz-Nle-Lys-Arg-Arg-AMC, Boc-Gly-Arg-Arg-AMC and Boc-Gly-Lys-Arg-AMC (Bachem AG, Bubendorf), which were dissolved in dimethyl sulfoxide for preparing stock solutions (100 mM). All enzymatic experiments were performed in triplicate and data are presented as mean ± SD, while IC_50_, Km and *K*i were obtained by fitting with GraphPad Prism 7.0 (39).

The pH dependence was measured with a protease concentration of 50 nM and substrate (Bz-nKRR-AMC) concentration of 250 μM at 0.5 pH intervals using the following buffers: 50 mM citrate-phosphate buffer for pH 4-5, 50 mM phosphate buffer for pH 5.5-8, 50 mM Tris-HCl buffer for pH 8.5-9.5, and 50 mM Na-bicarbonate buffer for pH 10-10.5.

For steady state kinetics, we used the exactly the same buffer as a previous one on profiling substrate specificity for the NS2B-NS3pro complexes of all four Dengue serotypes (17): 50 mM Tris-HCl at pH 8.5, except that no glycerol was added as it would lead to significant NMR signal broadening due to high viscosity. Briefly, Zika protease at 50 nM was incubated with substrates ranging from 10 to 1000 µM in 100 μl assay buffer at 37 °C. Progression of enzymatic reaction was monitored as an increase in fluorescence at λ_ex_ of 380 nm and λ_em_ of 450 nm. Fluorescence intensity is reported in arbitrary units. Initial fluorescence velocities (relative fluorescence units/sec) were calculated and curves were fitted to the Michaelis-Menten equation by nonlinear regression For IC_50_ determination, the protease at 50 nM was preincubated at 37 °C for 30 min with p-Nitrophenyl-p-guanidino benzoate at various concentrations and the reaction was initiated by adding Bz-nKRR-AMC to 250 μM. For *K*i determination, the assay was performed with different final concentrations of the inhibitor and substrate. The protease at 50 nM was incubated with the inhibitor at different concentrations for 30 min at 37 ºC. Subsequently, the reaction was initiated by addition of the corresponding concentration series of the substrate as described. The *K*i was obtained by fitting in the non-competitive inhibition mode (36).

### Homology modeling and molecular dynamics (MD) simulations

Protein sequences of the Zika NS2B and NS3pro were aligned with the corresponding sequences of the Dengue NS2B-NS3pro complex (12) and 30 homology models were built up with MODELLER 9v11 software package (26), and the selected model has the lowest discrete optimized protein energy (DOPE). Structural parameters of the Zika homology model and the Dengue NS2B-NS3pro complex (12) were very similar and thus judged to be suitable for MD simulation studies.

The protocol for the MD simulations of Zika and Dengue NS2B-NS3pro complexes in both open and closed conformations is exactly the same as we previously used for the Dengue NS2B-NS3pro complex in the open conformation (12). MD simulations of the closed conformation were performed on the crystal structures of Zika (5LC0) (25) and Dengue (3U1I) (31) NS2B-NS3pro complexes with their small molecule inhibitors removed. Four independent 100-ns MD simulations were conducted on Zika and Dengue NS2B-NS3pro complexes in both open and closed conformations.

The simulation cell is a dodecahedron with a periodicity of 12 Å between the protein and the box walls to ensure protein will not interact with its own periodic images. Water molecules using the TIP3P model are used to solvate the MD system. 8 Na^+^ ions were added into the Zika NS2B-NS3pro to make the charge of the respective MD system neutral. The solvated Zika homology NS2B-NS3pro complex was first energy minimized with 1000 steps of steepest descent using GROMACS version 4.5.7 (40) using AMBER all-atom force field (41). The time step was set as 2 fs. Prior to MD production simulation, two stages of equilibration were done independently for each MD simulation set. First equilibration phase was done in NVT ensemble for 100 ps with the following settings: long-range electrostatic interactions calculated with fast particle-mesh Ewald summation method; a cutoff of 9 Å for electrostatic interactions and a cutoff of 14 Å for van der Waals interactions; the temperature was kept constant at 300 K with Berendsen’s coupling. Second equilibration phase was done in NPT ensemble for 100 ps with the same settings as the first equilibration phase and some additional settings: pressure coupling was held at 1 bar with Parrinello-Rahma coupling; isothermal compressibility was set at 4.5*10 bar. Heavy atoms of proteins were position-contrained using LINCS algorithm (42). The MD production simulation was done with the same settings with NPT equilibration without the position-contraints on heavy atoms of protein and temperature coupling was done with Nose-Hoover.

## ACKNOWLEDGEMENT

This study is supported by Ministry of Education of Singapore (MOE) Tier 3 Grant R-154-002-580-112; Tier 2 Grant MOE2015-T2-1-111 and National Research Foundation of Singapore (NRF) R-154-002-529-281 to Jianxing Song. The funders had no role in study design, data collection and analysis, decision to publish, or preparation of the manuscript.

## AUTHOR CONTRIBUTIONS

Conceived the idea: JXS; Performed the experiments: AR LZL SS; Prepare figures and wrote the paper: JXS.

